# Pangenome evolution in *Escherichia coli* is sequence type, not phylogroup, specific

**DOI:** 10.1101/2022.05.20.492775

**Authors:** Elizabeth A. Cummins, Rebecca J. Hall, Chris Connor, James O. McInerney, Alan McNally

## Abstract

The *Escherichia coli* species contains a diverse set of sequence types and there remain important questions regarding differences in genetic content within this population that need to be addressed. Pangenomes are useful vehicles for studying gene content within sequence types. Here, we analyse 21 *E. coli* sequence type pangenomes using comparative pangenomics to identify variance in both pangenome structure and content. We present functional breakdowns of sequence type core genomes and identify sequence types that are enriched in metabolism, transcription and cell membrane biogenesis genes. We also uncover metabolism genes that have variable core classification depending on which allele is present. Our comparative pangenomics approach allows for detailed exploration of sequence type pangenomes within the context of the species. We show that pangenome evolution is independent of phylogenetic signal at the phylogroup level, which may be a consequence of distinct sequence type-specific driving factors relating to ecology and pathogenic phenotype.

**Data Summary:** Supporting data and code have been provided within the article or through Supplementary Data files available at https://doi.org/10.6084/m9.figshare.19793758. Custom Python scripts used to perform analyses are available at github.com/lillycummins/InterPangenome unless otherwise stated in the text.

## Introduction

*Escherichia coli* is a genotypically and phenotypically diverse species that inhabits a multitude of varying environments and is one of the most well-studied bacteria. The species is divided into eight main phylogroups: A, B1, B2, C, D, E, F, and G. Within phylogroups, further subdivision into clonal complexes and sequence types (STs) can be achieved by multilocus sequence typing (MLST) [1; 2]. The Warwick/Achtmann MLST scheme of *E. coli* is based on variations of seven house-keeping genes and has resulted in the generation of a vast multitude of STs. Coinciding with rich phenotypic heterogeneity, there is no gene pool barrier within *E. coli* [3] meaning that genetic material can be freely exchanged between pathogens and commensals. Therefore, some *E. coli* STs could conceivably act as vital genetic repositories in the development of important characteristics such as pathogenicity or antimicrobial resistance within other STs. The extent to which any given ST, or group of STs, acts as a reservoir (genetic source) or recipient (genetic sink) in the exchange of genetic information is currently not well known. The relatively recent development of pangenomics [4] provides a useful perspective we can use to interrogate the genetic contents of available genetic repositories provided by different *E. coli* STs and gain further insight into how these gene collections are structured and evolve.

The pangenome represents the set of all genes present in a given population [4; 5]. Pangenomic studies have been performed to understand *E. coli* at the species level [3; 6; 7; 8; 9] but comparative pangenomics analyses between STs within this species can potentially add to our understanding of the evolution of the species. The influence of population structure [8] and also the presence of complex epistatic relationships [10] are increasingly being acknowledged to have a major effect on the evolution of prokaryote pangenomes. Whelan and colleagues, for instance, noted that asymmetrical gene dependencies (e.g. the presence of *geneX* first requiring the presence of *geneY*, but not vice versa) cannot be uncovered by the consideration of coincident gene patterns alone. Conditional gene relationships can exist between genes, between sequence variants, or between genes and variants [11]. Inter-pangenome analysis -comparative analysis of closely-related pangenomes - provides an excellent mechanism for generating prioritised lists of putative dependencies between genes. Inter-pangenome analysis can show whether a gene’s classification as core in one ST is also core in a different ST. Interpangenome analysis can also assess differences in functional composition between closely-related pangenomes. The functional contents of a pangenome (whether species-level or ST-level) reflects the biological processes occurring within the given population, such as niche adaption [12; 13], or the evolution of important phenotypes, such as drug resistance [14].

An in-depth study of an ST131 pangenome revealed clade-specific diversity in colonisation and metabolism genes in the accessory genome of the globally dominant multidrug-resistant sub-lineage of ST131, clade C (H30Rx) [14]. The reported diversity was not due to the presence of unique genes, but rather the presence of unique alleles. Allelic diversity as a signature of selection has now also been observed in ST167 [15] and ST410 [16]. Allelic variation in metabolic genes has been described as an early warning sign of multidrug resistance with metabolic flexibility potentially being a key trait in multidrug resistant clones [17]. ST131 is one of the few *E. coli* STs to have undergone detailed pangenome analysis [14; 18; 19]. Understanding of *E. coli* STs on a comparative pangenome level is currently limited in terms of comparative analyses, with little known about how ST-level pangenome evolution is occurring. We wish to test the hypothesis that the different *E. coli* ST-level pangenomes do not evolve in the same way, by the gain and loss of the same kind of genes, but that their evolutionary histories and trajectories differ in significant and meaningful ways.

Here, using one of the biggest collections of *E. coli* genomes used to date, we further develop our understanding of *E. coli* pangenome dynamics and evolution by splitting *E. coli* into its constituent STs and comparing and contrasting the fates of these STs in the context of their respective pangenomes. We introduce an ST-focused approach to investigating evolutionary trends of pangenomic structure and contents, including the presence of sequence variants of metabolism genes, within E. coli. We addressed the following objectives: (i) to establish a conservative *E. coli* core genome, (ii) to assess whether ST pangenomes vssary in structure, (iii) to assess whether some ST pangenomes are enriched for specific biological processes, (iv) to assess the level of metabolic variation across ST pangenomes (given the potential link to multidrug resistance), and (v) to evaluate the potential for STs to act as genetic sources or sinks. We find that the distribution of genes across COG functional categories within an ST-core genome is not dictated by being in a given phylogroup and that enrichment occurs in specific functional categories that vary by ST. We also uncover conditional genetic relationships within core genomes, and we find that sequence variants differ in core classification within and between STs. Inter-pangenome analysis allows us to highlight how pangenome evolution is heterogeneous across a species and is independent of phylogeny, and we further our understanding of how collections of genes vary and evolve between STs.

## Methods

### Genome collection and sequence type pangenome analysis

20,577 publicly available *E. coli* assemblies were downloaded from EnteroBase [20] with a custom Python script (github.com/C-Connor/EnterobaseGenomeAssemblyDownload). Genome similarity was estimated using Mash [21] to ensure no duplicate entries were included in the dataset. The assemblies covered six phylogroups and 21 different STs of E. coli; ST3, ST10, ST11, ST12, ST14, ST17, ST21, ST28, ST38, ST69, ST73, ST95, ST117, ST127, ST131, ST141, ST144, ST167, ST372, ST410, ST648 (Table 1). Phylogroup G is an intermediate group between B2 and F that was characterised in 2019 by Clermont et al. [22]. This phylogroup is not included in the current analysis because it was unknown at the time of data collection.

**Table 1.** Summary of pangenome analyses. (Pangenome size does not include paralogs).

| Phylogroup | Sequence Type | No. of Genomes | No. of Core Genes | Pangenome Size | Core/Pan. (%) |
| --- | --- | --- | --- | --- | --- |
| A | ST10 | 2370 | 3066 | 27634 | 11.10 |
| A | ST167 | 115 | 3675 | 9035 | 40.68 |
| A | ST410 | 1006 | 3272 | 16223 | 20.17 |
| B1 | ST17 | 1884 | 4003 | 11870 | 33.72 |
| B1 | ST21 | 2411 | 4058 | 10671 | 38.03 |
| B1 | ST3 | 40 | 3749 | 7933 | 47.26 |
| B2 | ST12 | 283 | 3816 | 10531 | 36.24 |
| B2 | ST127 | 232 | 3891 | 9696 | 40.13 |
| B2 | ST131 | 3186 | 3460 | 15665 | 22.09 |
| B2 | ST14 | 62 | 3881 | 6825 | 56.86 |
| B2 | ST141 | 91 | 3879 | 8217 | 47.21 |
| B2 | ST144 | 65 | 3908 | 6938 | 56.33 |
| B2 | ST28 | 46 | 3630 | 7346 | 49.41 |
| B2 | ST372 | 54 | 3752 | 7514 | 49.93 |
| B2 | ST73 | 873 | 3789 | 11865 | 31.93 |
| B2 | ST95 | 758 | 3815 | 11933 | 31.97 |
| D | ST38 | 617 | 3780 | 14443 | 26.17 |
| D | ST69 | 696 | 3771 | 14808 | 25.47 |
| E | ST11 | 5137 | 3944 | 12904 | 30.56 |
| F | ST117 | 269 | 3722 | 11055 | 33.67 |
| F | ST648 | 382 | 3674 | 11610 | 31.65 |

Genes within each genome were annotated using Prokka (v1.12) [23]. Genomes were grouped by ST (using the EnteroBase classification) for individual ST-level pangenome analyses using Panaroo (v1.1.2) [24] with a 0.95 sequence identity threshold and a 0.99 core-genome sample threshold to allow the inclusion of unique core gene alleles in the accessory genome. We use the terms ‘gene cluster’ or ‘gene’ to refer to an orthologous gene group constructed by Panaroo. Linear regression was performed using Python scikit-learn (sklearn) LinearRegression module.

### Assignment of COG functional categories

The linear reference genome provided by Panaroo [24] for each ST pangenome was split into two lists of its respective core and accessory gene clusters. The nucleotide sequence for each gene cluster was translated using a custom Python script (github.com/C-Connor/GeneralTools) to obtain a protein sequence for each cluster. These protein sequences were used to characterise gene function. Gene clusters were assigned COG functional categories [25] using eggNOG-mapper (v2.0.8) bestOG assignment [26] and the eggNOG database [27] with sequence searches performed by DIAMOND (v2.0.7) [28]. Gene clusters that did not return a match within the eggNOG database were categorised under ‘?’. Heatmaps and clustermaps displaying distribution of COG categories across STs were made with seaborn (v0.11.2).

An ST was labelled as notably enriched in COG category ‘X’ if the percentage of ST-core genome designated to category ‘X’ lay above the upper quartile plus 1.5 times the inter-quartile range for all ST-core genomes in that category.

Functional domain annotation was performed with InterProScan (v5) [29; 30].

### Distribution of sequence type core genomes

Custom ABRicate databases were made for the core genome of each ST using the representative gene cluster nucleotide sequences from Panaroo and the --setupdb option in ABRicate (v0.8.7) (github.com/tseemann/abricate). The bottom 5th percentile of the average coverage distribution for each set of ST-core genes was removed to ensure any incorrectly called core genes were not included in our analyses. Mass screening across all 20,577 assemblies was carried out for each ST core database with ABRicate (21 searches in total). The results were summarised and partial hits (instances where a gene hit was split over multiple contigs) were accounted for and processed with a custom Python script. The average proportion of gene covered for each core gene cluster per ST was calculated.

### Core metabolic reconstructions

Metabolic models were constructed for the core genome of each ST using CarveMe [31]. Representative core gene cluster nucleotide sequences for each ST were used as input and the CarveMe algorithm was executed using the default settings. The number of metabolic reactions and metabolites in each ST core metabolic profile were counted using the Python COBRA package [32].

### Unique core metabolic reactions and genes

Metabolic reactions uniquely present in a single ST core metabolic reconstruction were extracted using a custom Python script. Unique reaction names were manually searched on the BiGG database website [33] to find the related gene names for each reaction. These related genes of interest (GOIs) were searched for in the descriptors of the ST-core gene sequences for the ST the related reaction was uniquely present in. These sequences were combined to construct a custom ABRicate database to perform a mass screening for the GOIs across all 20,577 assemblies with ABRicate. ABRicate results were summarised and processed using the same method previously described for the distribution of core genomes. A clustermap displaying the varying presence of the GOIs across 21 STs was made with seaborn (v0.11.2).

## Results

### A 2,172 gene cluster *E. coli* core genome

The size and content of the *E. coli* core genome has been estimated in previous studies [8; 34; 35; 36], but not explicitly using a collection of genomes as large as the dataset considered in this work. Here, we provide a representative *E. coli* core gene list. There were n = 2, 172 gene clusters identified that had a mean percentage coverage above 98% across all 20,577 assemblies. A list of these core genes and their nucleotide sequences are provided in File F1. Grouping these core genes by COG category showed that genes of unknown function (category S) were the largest functional group (18.7%). A breakdown of the functional composition of the core genes can be seen in Fig. 1. This large percentage of species-level core genes with unknown functions highlights that, despite extensive study and characterisation, there is still a great deal of information to be uncovered regarding core genes of E. coli.

**Fig. 1.**
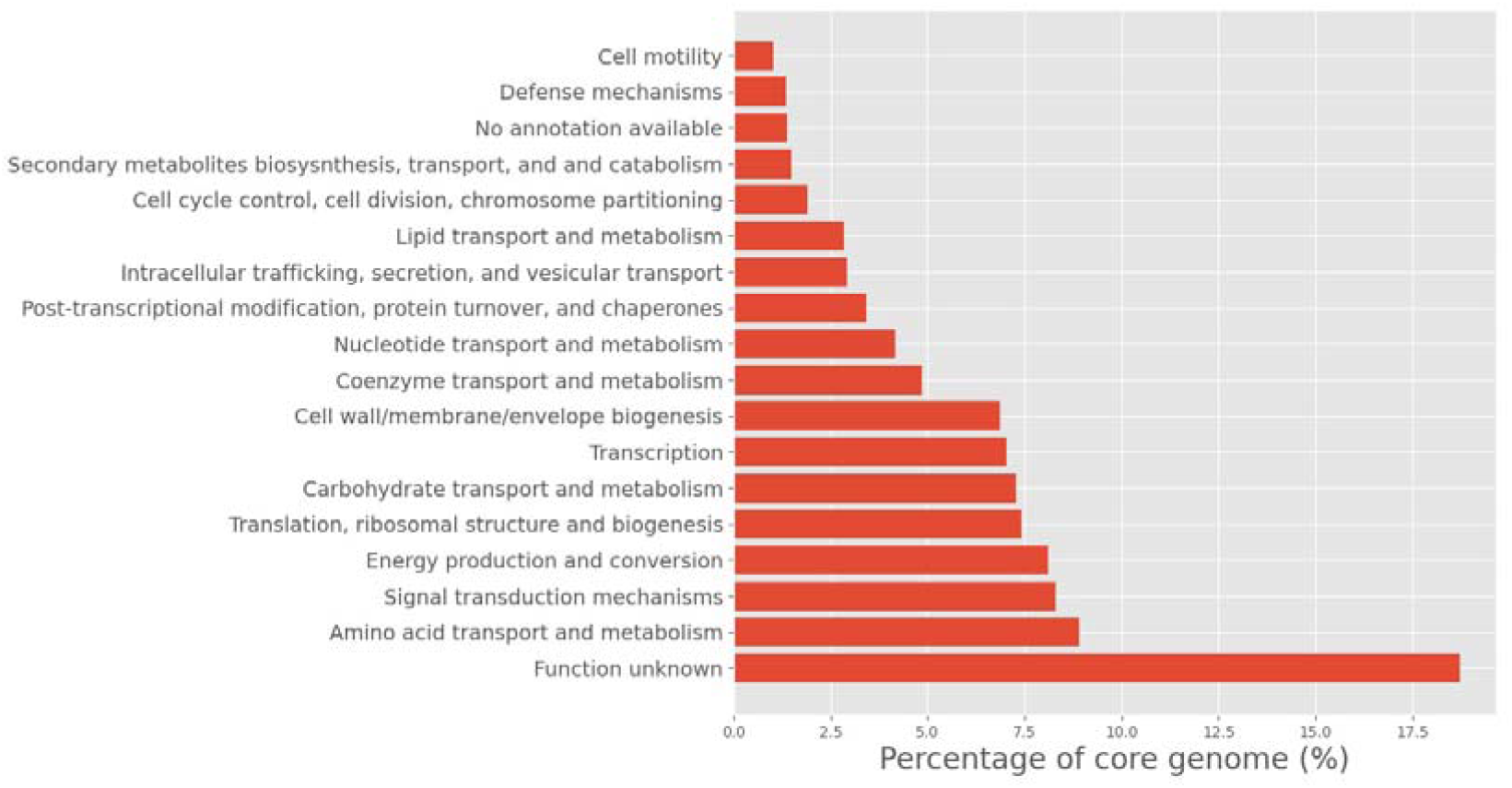
Functional breakdown of 2,172 core *E. coli* genes. Functional classes are based on COG categories.

### Pangenome structure varies between *E. coli* sequence types

To assess the level of variation between ST-level pangenomes we first considered variation in the context of structure. We assembled the pangenomes of 21 STs using Panaroo (v1.1.2). Pangenome sizes ranged from 6,825 to 27,634 gene clusters (Table 1), with an average size of 11,653 gene clusters and core genome sizes ranged from 3,066 to 4,058 clusters with an average of 3,738 clusters per ST pangenome. Neither core gene number (r^2^ = 0.005, Ordinary Least Squares), nor total gene number (r^2^ = 0.249, Ordinary Least Squares) were a function of ST sample size. Consequently, ST pangenomes exhibited variation in the core gene number as a percentage of the total pangenome size, which suggests that there is no uniform pangenome structure within *E. coli*. Variation in this percentage was highest for those STs with the fewest genomes, but even when the STs with greater than 100 genomes were considered, the variation in core gene number as a percentage of the total pangenome size extended from 11.1% (ST10) to 56.86% (ST14).

### Sequence type-specific core functions vary between sequence types

For the purposes of this study, we define three pangenome segments that are analysed and discussed throughout this work. Firstly, the ‘species core genome’ is the set of genes common to all genomes in this study. Next, the ‘non-species STX-core genome’ is the set of all genes considered core to STX, with the species core genome removed. Finally, the ‘STX-specific core genome’ is the set of genes that are found to be core only in STX and no other ST. These pangenome segments are conceptually visualised in Fig. 2. We calculated the percentage of each non-species ST-core genome that was assigned to each of the COG functional categories. As we were interested primarily in functional differences between STs, category S (function unknown) and ‘?’ (no functional annotation available) were masked from visualisation in Fig. 3 as they were always the largest two categories.

**Fig. 2.**
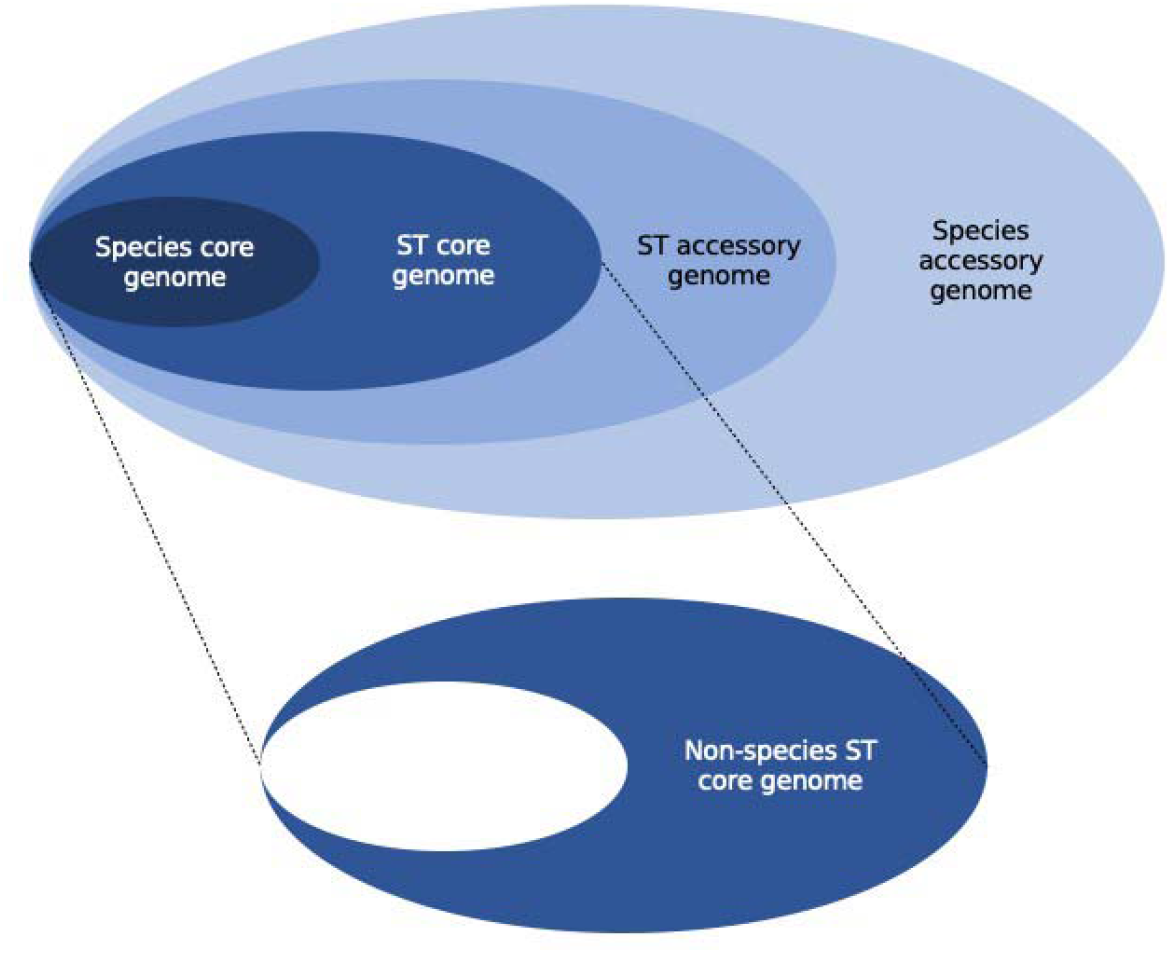
Depiction of pangenome segments used in this analysis.

**Fig. 3.**
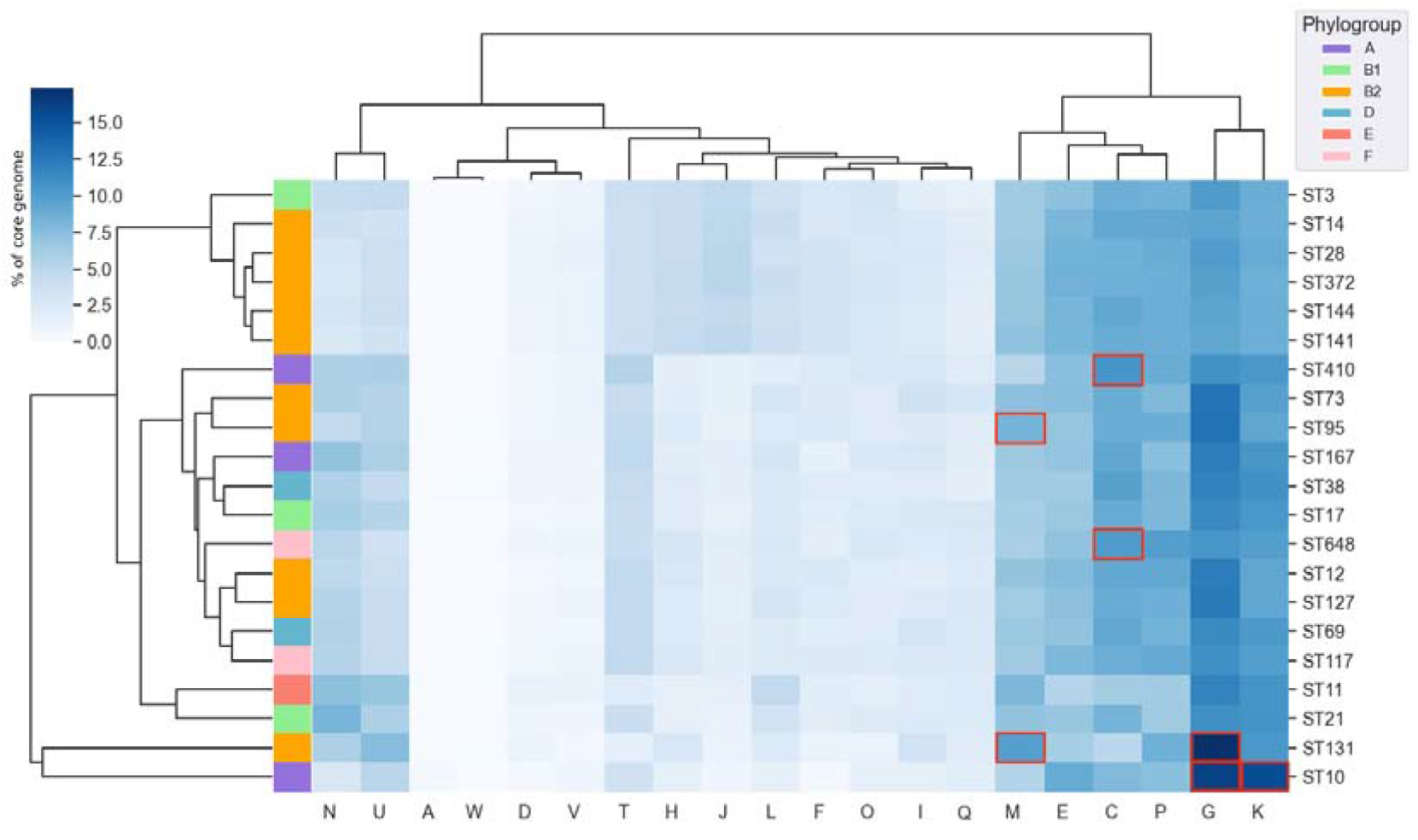
Hierarchically clustered (by percentage presence in core genome row-wise and column-wise) heatmap showing percentage of non-species ST core genomes classified into 20 functional COG categories. COG categories that are notably enriched in a sequence type are outlined in red. Functional COG categories: A - RNA processing and modification, C - energy production and conversion, D - cell cycle control and mitosis, Eamino acid metabolism and transport, F nucleotide metabolism and transport, G - carbohydrate metabolism and transport, H - coenzyme metabolism, I - lipid metabolism, J - translation, ribosomal structure and biogenesis, K - transcription, L - replication and repair, M - cell wall/membrane/envelope biogenesis, N - cell motility, Opost-translational modification, protein turnover, chaperone functions, P - inorganic ion transport and metabolism, Q - secondary metabolites biosynthesis, transport and catabolism, T - signal transduction, U - intracellular trafficking and secretion, V defence mechanisms, W - extracellular structures.

Hierarchical clustering of the percentage of each ST-core genome assigned to 20 COG functional categories highlighted the ST131 and ST10 non-species core genomes as having the most distinct functional profiles (Fig. 3). The accessory genomes were also functionally categorised, however in all ST pangenomes the accessory genome was dominated by genes of unknown function (data not shown). The data show that ST pangenomes do not possess uniform core functional profiles and additionally, this observed variation is not heavily influenced by the identity of the phylogroup.

We also examined variation in non-species ST-core genomes, and in particular their propensity to be differentially enriched in specific biological processes of a particular function. ST95, ST410, ST648, ST131, and ST10 exhibited notable functional enrichment in four COG categories in their non-species ST-core genomes (see Methods). ST95 was notably enriched in genes linked to cell membrane biogenesis (category M); ST10 was notably enriched in genes pertaining to transcription (category K) and carbohydrate metabolism and transport (category G); ST410 and ST648 were notably enriched in energy production and conversion genes (category C); and ST131 was notably enriched in genes pertaining to cell membrane biogenesis (category M) and carbohydrate metabolism and transport (category G). These enriched categories are highlighted in Fig. 3. This suggests that genes encoding these functions may be particularly influential in these STs.

### ST131 and ST10 pangenomes possess multiple alleles related to carbohydrate metabolism and transport

We have identified two non-species ST-core genomes, ST10 and ST131, that are notably enriched in carbohydrate metabolism and transport (category G) genes. To investigate whether this enrichment was related to metabolic diversity, we explored the presence of alleles within the category G genes in the ST131 and ST10 pangenomes. The non-species ST131-core genome includes n = 100 gene clusters linked to carbohydrate metabolism and transport (category G), of which 64% are indicated to have multiple alleles present, as different gene clusters, in the ST131 pangenome (File F2). These include, but are not limited to, *manRXZ, sorABFM, malPX* and *gatABCYZ*, involved in the mannose, sorbose, maltose, and galactose phosphotranserfase systems [37; 38; 39; 40]. The non-species ST10 core genes in category G (File F3) that have multiple alleles present across the ST10 pangenome include *mngAB*, involved in mannose transport and metabolism [41] and sugar efflux transporters setAC [42]. A full summary of the number of genes present as multiple alleles per enriched COG category is provided in File F4. Beyond multiple alleles of carbohydrate metabolism genes being present across the pangenome, certain genes were present as multiple alleles within the ST131 core genome. The non-species ST131-core genome possessed two alleles of each of the following genes: *fruA, gatC, kdgK, nagB, tabA, and uxaA*. Multiple alleles of carbohydrate metabolism genes were not present within the non-species ST10-core genome, but there were multiple alleles present of three transcription genes; *glpB, hcaR, and mngR*.

Further investigation revealed that the fructose phosphotransferase system (PTS) gene, *fruA*, beyond being present as two alleles in the non-species ST131-core genome (as clusters ‘fruA_2’ and ‘fruA_3_fruA_1’) was in fact present as four gene clusters across the ST131 core genome; ‘fruA_1’ and ‘manP_fruA_4_fruA_1’ clusters were found to be present in the species core genome. To investigate the functionality of these four gene clusters, functional domain analysis was carried out using InterProScan [29]. Three distinct functional domains corresponding to the PTS system EIIA, EIIB, and EIIC components [43] were identified within the four *fruA* clusters. ‘fruA_1’ possessed both EIIB and C, whilst ‘manP_fruA_4_fruA_1’ and ‘fruA_3_fruA_1’ possessed only EIIC (Fig. 4). ‘fruA_2’ encodes the PTS EIIA component that is associated with *fruB* rather than *fruA* [44]. This discrepancy is likely attributable to the Panaroo cluster naming algorithm rather than being of biological significance. The presence of these four gene clusters is shown in Fig. 4. The three correctly named *fruA* clusters are distinct (greater than 5% sequence divergence), highly conserved, seemingly functional, and encode non-truncated peptide sequences which suggests annotation error, degradation, or pseudogenisation are unlikely to be responsible for this multiplicity.

**Fig. 4.**
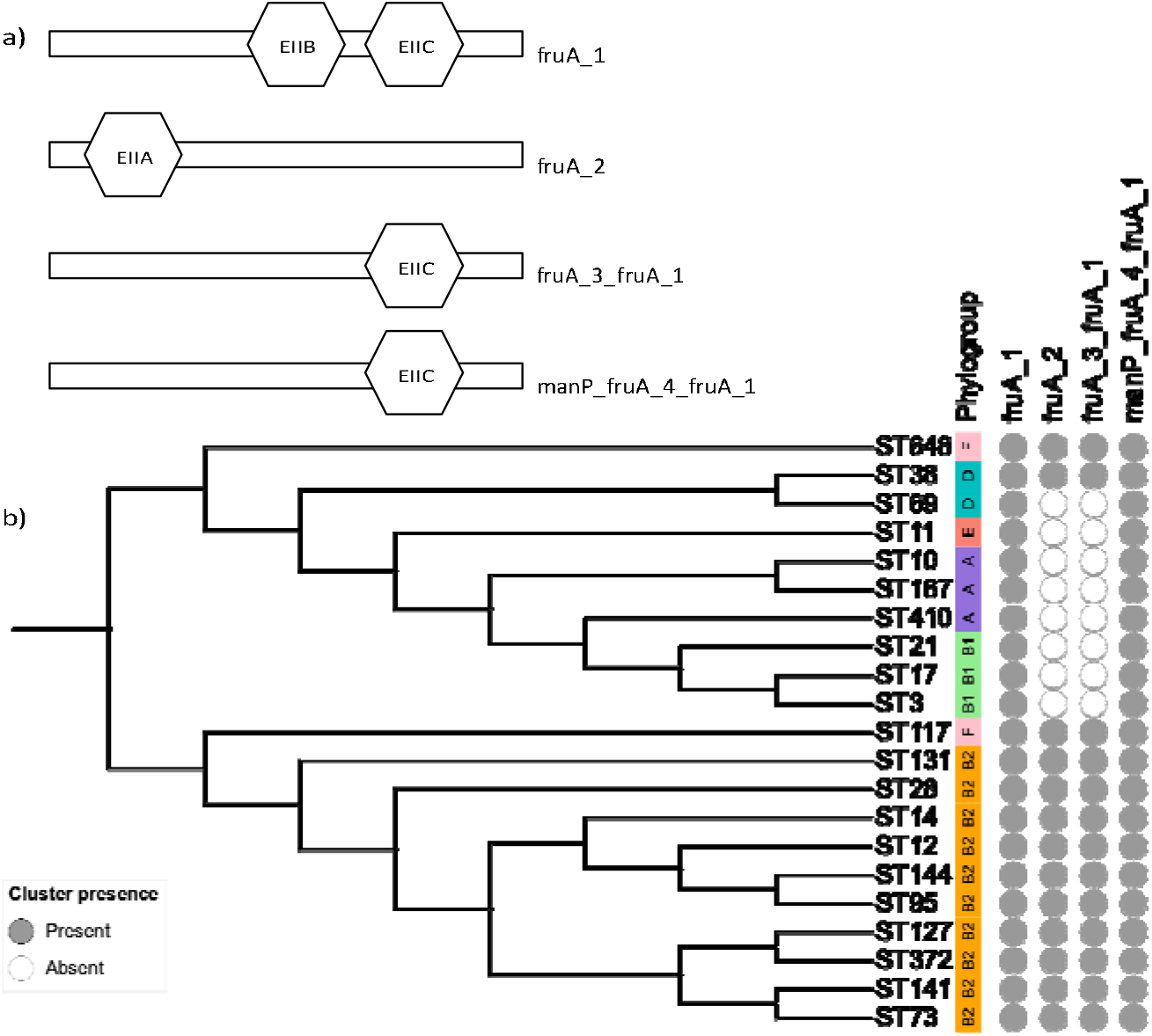
a) Phosphotransferase system components encoded by four fruA annotated clusters and b) cluster presence across an *E. coli* phylogeny, created using iTOL [45].

### The ST131 pangenome has distinct gene presence patterns at phylogroup and species level

Further investigation of the presence patterns of genes from ST pangenomes enriched in specific COG functional categories (outlined in Fig. 3) across other ST pangenomes revealed that the ST131 pangenome displays gene presence and absence patterns that are distinct from other phylogroup B2 ST pangenomes. The ‘mhpA’ gene cluster from the ST410 pangenome, or ‘mhpA_1’ gene cluster from the ST648 pangenome, is present in only the ST131 pangenome out of the ten B2 phylogroup ST pangenomes (Fig. S1, S2). *mhpA* encodes a 3-(3-hydroxyphenyl)propionate hydroxylase involved in phenylalanine metabolism [46]. Similarly, the ‘mngA_1’, ‘mngB’ (Fig. S3) and ‘mngR_1’ (Fig. S4) clusters from the ST10 pangenome are only present in the ST131 pangenome out of all B2 phylogroup ST pangenomes. There are also gene clusters (‘rspA_1’, ‘hbp’, ‘tsx_1’, ‘fimC_1’) from the ST95 pangenome that are absent in only the ST131 pangenome out of the B2 phylogroup ST pangenomes (Fig. S5).

The ST131 pangenome possesses core genes that are not seen in any other *E. coli* ST pangenome considered in this study. Notable presence patterns within ST131’s enriched carbohydrate transport and metabolism core genes (Fig. S6) include the uniquely present ‘group_3501’ and ‘yihP_yicJ_3_yicJ_1’ gene clusters. These clusters were not detected in any other ST pangenome. Functional annotation of ‘group_3501’ suggests this gene encodes a glycosyl hydrolase and eggNOG provided xylS as an annotation for this gene cluster. In the KEGG orthology database [47; 48], *xylS* is synonymous with *yicI* [49]. With this connection, we postulate that these two gene clusters, uniquely present in the ST131 pangenome, are involved in the same xyloside metabolic pathway. The nucleotide sequence was searched against the uniprot [50] database resulting in a top hit of 85.3% similarity to a putative glycosyl hydrolase from *Citrobacter rodentium and Citrobacter freundii*.

### Alleles of metabolism genes vary in core status across sequence types

To examine metabolic diversity within ST-core genomes in more detail, we performed metabolic reconstructions for the core genome of each ST pangenome using CarveMe [31] to create a ‘ST core metabolic profile’ so that gene information could be extrapolated to utilisation and specific metabolic pathways. Comparison of the 21 ST core metabolic profiles uncovered 825 metabolic reactions that were found in at least one ST core metabolic profile, but not common to all STs (*i*.*e*. not a species-core reaction). We focused further analysis on metabolic reactions that were uniquely present in a single ST core metabolic profile. Tracing uniquely present reactions within the BiGG database [33] back to their related gene names, and then searching for these names within our dataset, led us to a subset of gene clusters with non-ubiquitous presence patterns (Fig. 5). These selected clusters were *fhuA* (iron acquisition) [51], *pduCDEF* (propanediol utilisation) [52], *mntH* (manganese transport) [53], and the hydratase *crt* [54].

**Fig. 5.**
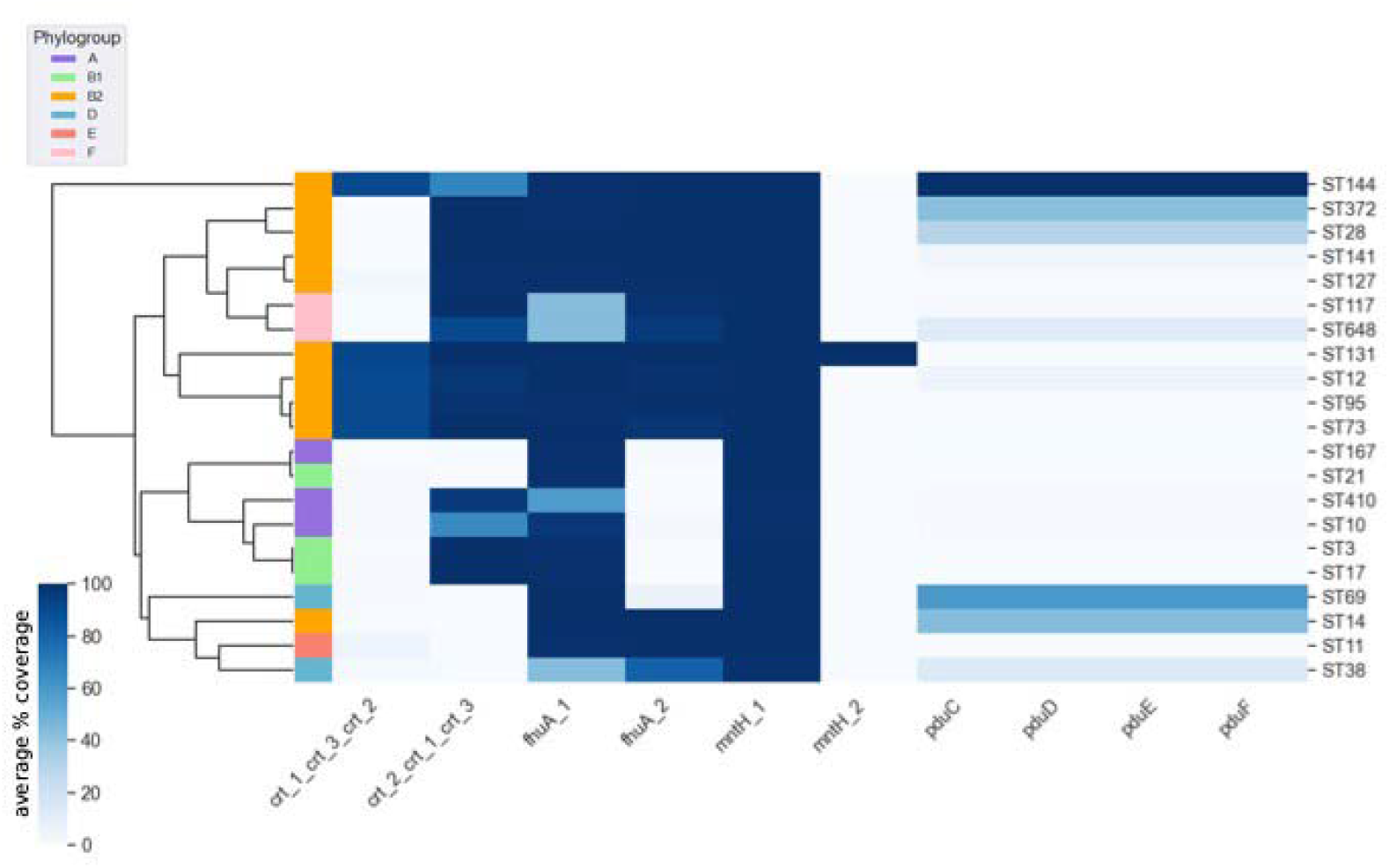
Hierarchical row-wise clustering of the average presence of *crt, fhuA, mntH* and *pduCDEF* gene clusters across 21 sequence types of *E. coli*.

The manganese transporter *mntH* [53] has two alleles present in the core genome of ST131 (Fig. 5), which raises the question of why there is a fixed second allele and also why this has not happened in another ST pangenome. Functional domain analysis of ‘mntH_1’ and ‘mntH_2’ returned the same InterPro annotation accession number for both alleles. Similarly, *fhuA* and *crt* have two alleles simultaneously present in the core genomes of multiple STs (Fig. 5). The crt alleles are also involved in a conditional relationship. From the clustermap in Fig. 5, we see that the ‘crt_1_crt_3_crt_2’ cluster is only present in an ST pangenome when ‘crt_2_crt_1_crt_3’ is also present, with the possible exception of ST144. Additionally, the ‘fhuA_2’ cluster is only present, excluding phylogroup F STs, when ‘fhuA_1’ is also present in the ST, with the possible exception of ST38.

Fixation of the propanediol utilisation operon *pduCDEF* has occurred uniquely in ST144 (Fig. 5). These genes are reported in other STs (ST372, ST28, ST141, ST648, ST12, ST69, ST14, ST38) at lower average frequencies, showing that *pduCDEF* are accessory genes intermittently present within these STs. The *pdu* operon is involved in anaerobic respiration which is used by enteropathogenic *Enterobacteriaceae* to out-compete existing intestinal microbiota during infection and is frequently reported in *Yersinia enterocolitica* and *Salmonella Typhimurium* [55]. However, this was considered to be a rare phenotype in *E. coli* [6]. Each of the four genes presented here provide additional evidence for sequence level variation in metabolism between *E. coli* STs.

### ST10 has the potential to be a genetic source for other *E. coli* sequence types

Evidence for a phylogroup or ST acting as a genetic source for other *E. coli* may arise in the form of an ST pangenome possessing a low (or no) amount of ST-specific core genes (genes that are classified as core in only one specific ST). To this end, gene clusters that were uniquely core to a specific phylogroup were examined first. The three ST-pangenomes in phylogroup A had no unique core genes (Table 2). Alleles including those of flagellar genes *fliDS* are present amongst the 52 B1 unique core genes. B2 ST pangenomes possessed 99 uniquely core gene clusters, including alleles of genes involved in central metabolism, *sucABCD*, fructose metabolism, *fruA*, and the decarboxylase *tabA*. The 14 unique core genes of the two phylogroup D ST pangenomes included alleles of a putative fimbrial protein, *yadNMV*. Extending this analysis from phylogroup to ST, we also considered unique ST-specific core genes. The number of alleles uniquely core to a single ST varied within and between phylogroups, with ST14 (phylogroup B2, n = 83) and ST21 (phylogroup B1, n = 74) encoding the largest number of unique core genes (Table 2). Whilst phylogroup A has no unique core genes, within this phylogroup only the ST10 pangenome had no reported unique ST-specific core genes; the ST167 and ST410 pangenomes were found to have 28 and six unique core genes, respectively. The ubiquity of the ST10 core genome across all other STs may be an indicator that this ST is likely to be capable of acting as a genetic source within E. coli.

**Table 2.** Numbers of genes uniquely core to a single phylogroup and sequence type.

| Phylogroup | Unique phylogroup-core genes | ST | Unique ST-core genes |
| --- | --- | --- | --- |
| A | 0 | ST10 | 0 |
|  |  | ST167 | 28 |
|  |  | ST410 | 6 |
| B1 | 52 | ST3 | 68 |
|  |  | ST17 | 14 |
|  |  | ST21 | 74 |
| B2 | 99 | ST131 | 21 |
|  |  | ST28 | 44 |
|  |  | ST14 | 83 |
|  |  | ST12 | 10 |
|  |  | ST95 | 8 |
|  |  | ST144 | 56 |
|  |  | ST141 | 38 |
|  |  | ST73 | 26 |
|  |  | ST127 | 14 |
|  |  | ST372 | 23 |
| D | 14 | ST38 | 23 |
|  |  | ST69 | 43 |
| E | 99 | ST11 | 99 |
| F | 1 | ST117 | 24 |
|  |  | ST648 | 40 |

## Discussion

Extensive phenotypic variation and the existence of diverse STs within *E. coli* is well documented [6; 56]. However, little is known about how the genetic repertoire of each ST varies in terms of pangenome structure and content, and consequently which genes are given core status within different ST pangenomes. We build upon previous work analysing a single *E. coli* species pangenome [6; 7] or *E. coli* ST pangenome [14; 18; 19] by performing large scale comparative analysis on 21 ST pangenomes constructed from over 20,000 genomes. We introduce the concept of comparative pangenomics with a method that interrogates ST pangenome content and structure variation across the species. We also classified the non-species ST-core genome of each ST pangenome into COG functional groups. Our study revealed variation in pangenome structure and core genome functionality both across and within *E. coli* phylogroups.

Previous estimates of the size of the *E. coli* core genome fell in the range of 1,000 to 3,000 gene clusters and were extrapolated from small genome collections ranging from 14 to 186 isolates [34; 35; 36]. We build upon this earlier work by estimating an *E. coli* core genome with a larger dataset. The non-uniformity in structure between *E. coli* ST pangenomes demonstrates the extent of the flexibility within this species and is a valuable lesson gained from comparative pangenomics. We also found that the function of genes given core status within an ST pangenome (the non-species ST-core genome) varied between STs, with certain ST pangenomes having notably higher percentages of genes in four functional COG categories: energy production, carbohydrate metabolism, transcription, and cell membrane biogenesis. This enrichment may signpost ST-specific adaptive evolutionary processes as a signature of selection via accumulation of allelic diversity. It is already known that there are large ecological variations in E. coli; isolates have been found as gut commensals in most animals, as well as in environmental samples, and can exist on a spectrum of pathogenicity ranging from complete commensal to strict pathogen. Our findings strengthen the concept that evolution towards a given ecology and pathogenic phenotype occur at an ST-by-ST level in E. coli.

Going beyond consideration of ST-core genomes as functional units, we attained more nuanced findings regarding ST-core gene variants. We found possession of multiple variants of carbohydrate metabolism genes in ST pangenomes. More diverse genes relating to metabolism, including clone specific SNPs in anaerobic metabolism loci within ST410 [16] and ST131 [14], have been reported previously. In these cases, the sequence diversity was attributed to differential evolution whereby selection for a process (enhanced anaerobic metabolism capabilities) rather than selection for a gene was occurring. Metabolic flexibility has also been proposed as a precursory stage to multidrug resistance [17]. We could conceivably extrapolate our findings, such as our observed diversity in a fructose metabolism gene within an ST-core genome, as a potential signature of an evolutionary selection pressure.

The fixation of the pdu operon in the ST144 pangenome suggests a unique evolutionary history. ST144 is a uropathogenic *E. coli* that shares the closest common ancestor with ST95 [57]. 1,2-propanediol is enriched in the mucosal lining of the intestine so the ability to utilise this alternative carbon source is advantageous in an inflamed gut [58]. Similarly, the ST131 pangenome has a second *mntH* allele and a glyosyl hydrolase linked to xyloside metabolism, among other distinct gene presence patterns which suggests a separate evolutionary trajectory for this ST. Recent mash-based analysis by Abram and colleagues has demonstrated notable differences between ST131 and other B2 strains that was significant enough to classify ST131 within the subgroup B2-1, said to have recently emerged from B2-2 [59]. The ability to discriminate between ST131 and the rest of the B2 phylogroup was attributed to the differential, rapid uptake of unique virulence factors and mobile genetic elements by ST131 [60]. The unique gene presence patterns we reported within the ST131 pangenome are consistent with this previous study [59].

Pangenomes can reflect the ecology of an organism [61; 62] so insight may be gained by translating gene presence/absence to, for example, niche occupation. Genes core to an ST, or a group of STs, provide indicators of evolutionary advantages in certain ecological settings and genetic backgrounds [63]. From our dataset, ST10 was the only ST pangenome to have no unique core genes. This weak unique core signature may reflect the heterogeneous nature of ST10 [59]. This aligns with previous work that has identified ST10 as a generalist lineage and a potential genetic reservoir for other *E. coli* lineages [6; 8].

We propose that selection and pangenomic evolution is independent of phylogroup phylogeny in E. coli. Our goal was to test whether *E. coli* ST pangenomes were evolving in a uniform way. The test of this hypothesis yields different perspectives if the comparison is carried out at the level of phylogroup or the level of ST. The variation in core functions between ST pangenomes is a clear signal of ST-specificity and we have shown throughout this work that ST pangenomes are distinct in different ways, from structure to alleles of genes varying in core status across ST pangenomes. We have also provided a putative list of core gene clusters from a dataset of over 20,000 *E. coli* genomes. The future of microbial genomics is in population genomics, and specifically comparative pangenomics. This dataset provides a wealth of information on *E. coli* as a species and variation across ST pangenomes.

## Author Statements

### Authors and contributors

EAC: methodology, formal analysis, visualisation, writing - original draft preparation and review and editing. RJH: writing – original draft preparation and review and editing. CC: data curation, writing – review and editing. JOM: conceptualisation, writing – review and editing. AM: conceptualisation, supervision, writing – review and editing.

## Acknowledgements

We would like to thank Dr. Steven Dunn for fruitful discussions on the visualisation process used in the work presented here.

